# Computational insights on the competition between electrotaxis and durotaxis

**DOI:** 10.64898/2026.02.02.700142

**Authors:** Pablo Sáez, Shardool Kulkarni, Custodio Nunes, Min Zhao, Elias Barriga

## Abstract

Understanding how cells migrate in response to external cues has important implications for biology, medicine, and bioengineering. Chemical, mechanical, and electrical signals are the primary drivers of directed cell migration, and each has been extensively studied over the past decades. Among them, chemical cues were the first to be investigated and remain the most widely studied due to their undeniable role in in vivo guidance. Mechanical signals—particularly substrate stiffness gradients—have gained prominence for their ubiquity across cell types and their potential to direct migration. More recently, growing evidence suggests that electrotaxis offers a highly precise and programmable means to guide cell movement. Despite this, these cues are often studied in isolation, whereas in vivo they typically coexist and interact. Using well-established biophysical models, we investigate how mechanical and electrical signals cooperate and how they can be engineered to compete for control over cell migration. We demonstrate that an electric field can override and even reverse durotaxis, with outcomes that depend strongly on the specific cell type. To address this large variability in controlling cell migration, we propose particular steps toward further exploration. To support such future research, we provide a freely available platform for predicting electro-mechanical interactions in cell migration, based on a given cell’s sensing and signaling characteristics, which could tailor the mechanical and electrical signals that arise naturally during organ development, cancer invasion, or tissue regeneration.

## Introduction

Cell migration is a fundamental biological process that underpins critical aspects of life, including embryonic development, immune responses, wound healing, and pathological conditions such as tumor invasion [1, 2, 3]. In both in vivo and in vitro settings, cells respond to various external cues to navigate their environment [4]. Moreover, the capacity to guide cellular movement using external stimuli is gaining traction in medicine—aiming to halt tumor invasion, enhance tissue repair, or accelerate wound healing [5, 1, 6]. In tissue engineering, this control over migration supports the spatial arrangement of cells within engineered constructs, enabling the formation of biomimetic tissues with functional architectures [7, 8].

Chemotaxis is among the most well-studied mechanisms governing directed migration, where cells move in response to gradients of soluble biochemical signals [9]. These gradients activate membrane-bound receptors and initiate downstream signaling cascades. Central to this process are the Rho family of GTPases, including RhoA, Rac1, and Cdc42, which regulate myosin and actin cytoskeleton dynamics, as the key drivers of the forces underlying cell migration [10, 11]. Specifically, RhoA promotes actomyosin contractility [12, 13, 14], Rac1 drives lamellipodia formation [15, 16, 17, 18]—aiding in spreading—and Cdc42 controls filopodia formation, essential for environmental sensing and directional probing [19, 20, 21]. Cross-talk among these GTPases allows for the spatial coordination of adhesion dynamics, with Rac1 and Cdc42 activity concentrated at the leading edge and RhoA predominantly active at the rear of migrating cells [22, 23].

In addition to biochemical cues, mechanical signals are now well recognized as pivotal regulators of cell migration. For instance, cells migrate along gradients of substrate stiffness and adhesion ligand density—phenomena known as durotaxis and haptotaxis, respectively [24, 25, 26]. These processes involve asymmetric adhesion forces. Similarly, during frictiotaxis—the migration toward regions of higher extracellular friction—mechanical friction is transduced into intracellular cues that induce polarization of the actomyosin network in integrin-independent migration [27]. Notably, effective friction has been shown to underlie both positive and negative durotaxis, suggesting that friction alone, whether in mesenchymal or amoeboid migration, can drive directed cell movement [28]. Although this friction differs in nature, we will hereafter refer to “durotaxis” to encompass the effective friction transmitted to the retrograde actin flow.

While often approached from a mechanical perspective, where cell motility is explained by the balance of forces alone [29, 30, 31, 28], durotaxis also engages intracellular signaling pathways similar to those activated in chemotaxis, which directly modify crucial motile forces of the cell for both homeostasis and pathological conditions. Integrin activation, for example, stimulates Rho GTPases, coordinating the organization of adhesion complexes and cytoskeletal remodeling [10, 11] and enabling cells to sense and adapt to mechanical forces.

Electrotaxis—the directed migration of cells in response to electric fields (EFs)—is another guidance mechanism observed in physiological contexts such as embryonic development [32] and wound healing [33], and it is present across a wide range of cell types [34, 35, 36, 37], and even whole tissues [38, 39]. Charged Membrane Proteins (CMPs) such as Epidermal Growth Factor Receptor (EGFR), membrane lipids, and Vascular Endothelial Growth Factor (VEGF) receptors, among others, have been shown to polarize during electrotaxis. Moreover, some of these polarized components may contribute to the asymmetric activation of Rho GTPases, aligning electrotaxis with the same intracellular machinery involved in chemo-taxis and durotaxis. Indeed, one of the most fundamental questions in electrotaxis is what specific CMPs are actually responsible for the downstream polarization of GTPases.

In summary, chemotaxis, durotaxis, and electrotaxis are all orchestrated through the inside-out and outside-in signaling pathways mediated by Rho GTPases. These directed migration mechanisms occur naturally in vivo and have been extensively studied in isolation. However, in physiological contexts, these cues often coexist, or one can be imposed artificially over the others to foster or arrest their effect in cell migration. Despite this, there is a surprising lack of research exploring how these extracellular signals interact synergistically to regulate cell migration, limiting the potential to efficiently control cell migration.

In this work, we investigate the competition between mechanical and electrical signals, as easily controllable cues, in directing cell migration. We use well established computational models to describe and integrate the mechano-chemical phenomena leading to each of these tactic cues. Our aim is to establish a hypothesis for how these cues compete to influence the underlying signaling networks and the cell’s motile machinery. To support further research and practical applications, we also present an open-access online platform (available on MATLAB Central File Exchange) that, given the specific characteristics of a cell line, predicts its migratory behavior—including direction, velocity, and intracellular dynamics of Rho GTPases and actomyosin structures.

## Results

To investigate how simultaneous mechanical and electrical stimulation influences cell migration, we adapt a minimal active gel model. Active gel models characterize the dynamics of the actomyosin network by solving the transport equations for actomyosin density in conjunction with the momentum balance equation for the retrograde flow [40, 41]. In short (see SI for details), the model computes the retrograde velocity *v* and the polymerization velocity against the cell membrane *v*^*p*^, from which we calculate the motion of the cell boundaries as a combination of inward retrograde flow and outward protrusion such that the velocities of the front and rear edges are given by 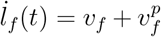 and 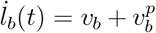, respectively. Then, the migration velocity of the cell is given as *v* = (*ŀ*_*f*_ (*t*) + *ŀ*_*b*_(*t*))/2 (see Fig. 1a).

**Figure 1.**
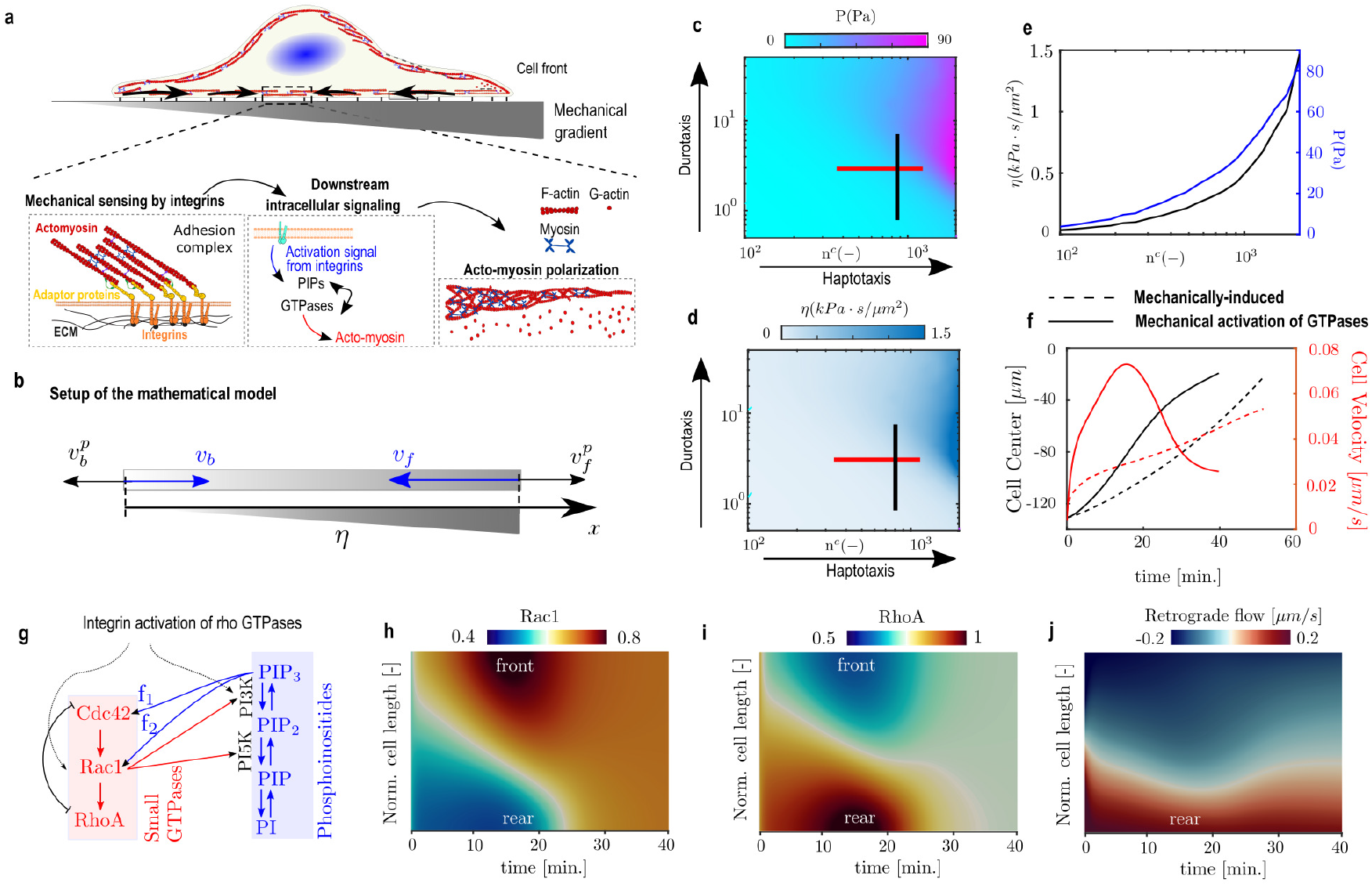
Durotaxis in single-cell migration. (a) Schematic of the main mechanisms involved in durotaxis, including integrin activation, downstream signaling, and actomyosin polarization. (b) Key variables in the physical model: retrograde flow velocity *v* (blue), indicated at the front and rear of the cell (subscripts *f* and *r*, respectively), and actin polymerization velocity (black). (c-d) Traction forces (c) and corresponding effective friction (d) in response to gradients in ECM stiffness and anchoring point density. Red and black lines represent traction and friction gradients of similar magnitude for haptotaxis and durotaxis, respectively. (e) A representative traction and friction profile used in the simulations. (f) Time evolution of cell center position (black line) and migration velocity (red line) in two conditions: (1) when durotaxis is driven solely by a friction gradient (dashed lines), as described in [28], and (2) when integrin activation also triggers downstream activation of PI3K and subsequently Rho GTPases. Incorporating biochemical signaling results in faster cell migration, due to increased actin polymerization at the front and enhanced retrograde flow at the rear of the cell. (g) Schematic of the Rho GTPase signaling network included in the mathematical model. Arrows indicate positive feedback (activation) from one component to another and tails indicate negative feedback (inactivation) of one component by another. *f*_1_ and *f*_2_ are the magnitudes of feedback for activating Cdc42 and Rac1 by PIP3. (h-j) Kymographs of variables along the normalized cell length (Y-axis) over time (X-axis) when mechanical activation of GTPases is included in the durotactic model: Rac1 (h), RhoA (i), and retrograde flow velocity (j). In all kymographs, the positions of the cell front and rear are indicated.

If no symmetry-breaking is induced, the cell will show symmetric flows and actomyosin densities, and, consequently, cell spreading [42, 43, 44]. This non-migrating, static cell mode has been reproduced using such active gel models [40] and also used to describe the transmission of forces between different cellular networks and adhesion structures [45].

During durotaxis, gradients in mechanical cues of the ECM induce asymmetric expression of cell adhesions, highly dynamic structures primarily mediated by transmembrane receptors such as integrins, which play a crucial role in guiding directed migration. The clutch model [46] has been instrumental in explaining and predicting cell adhesion behavior across various ECM conditions, including elastic [47, 48], viscoelastic [49, 50], and ligand density [51] (see [52] for a detailed overview). By coupling an active gel model of cell migration with a stochastic clutch framework, it is possible to describe how the active forces of the cell are transmitted asymmetrically outside the cell, inducing cell movement. This mathematical approach has been used, for example, to describe both positive and negative durotaxis across different cell types [28]. To develop our current model, we begin with a simplified version of this previous durotactic model [28].

First, we compute the traction forces using the clutch model, along with the effective friction (see SI) resulting from haptotactic and durotactic cues (Fig. 1c-d). We then extract the traction and friction profiles corresponding to specific haptotactic and durotactic gradients (represented by the red and black lines in Fig. 1c-d), and impose these gradients over a substrate of 250 *μ*m in length. Therefore, rather than dynamically solving the clutch model over time, we apply the precomputed effective friction along each position of the cell–substrate interface. To initialize our simulations, we assumed a cell with constant normalized acto-myosin density, zero retrograde flow, and placed the cell center at -125 *μ*m As previously demonstrated, cells migrate along friction gradients (Fig. 1f) [28], rather than stiffness gradients alone—challenging earlier assumptions and offering a theoretical rationale for previously observed phenomena described as positive and negative durotaxis. However, this durotactic model predicts a lower migration velocity compared to experimental observations [53, 54, 55]. One possible explanation is that the model accounts solely for the mechanical interactions mediated by cell adhesions, while neglecting the fact that, as discussed above, integrin activation also initiates Rho GTPase signaling, which could enhance the cell’s migration speed.

Integrin activation involves a conformational transition from a low-affinity (inactive) state to a high-affinity (active) state, regulated by intracellular adaptor proteins such as talin and kindlin. This process is controlled by bidirectional signaling: intracellular cues regulate integrin activation (inside-out signaling), while integrin–ECM interactions transmit signals back into the cell (outside-in signaling), influencing cytoskeletal organization and downstream signaling pathways. Notably, integrin engagement activates PI3K [56, 57, 58, 59]. In fibroblasts, adhesion to fibronectin has also been shown to promote transient Rac activation and sustained Rho activation [60, 61, 62, 63, 64]. Other studies have reported that integrin-mediated adhesion activates both Cdc42 and Rac1 [65, 62], while some suggest that Cdc42 is activated first and subsequently triggers Rac activation [63].

To incorporate these activation mechanisms, we coupled our durotactic model with a well-established Rho GTPase signaling model [66, 67, 68] This model includes both active and inactive forms of Rac1, Cdc42, and RhoA, as well as key phosphoinositides: PIP (*P*_1_), PIP2 (*P*_2_), and PIP3 (*P*_3_) (see Fig. 1g). To account for the activation of Rho GTPases by integrins, we follow a previous model [69] and consider that the conversion of PIP2 to PIP3 via PI3K depends on integrin activation, which we assume to be proportional to friction. Following experimental insights [56, 57, 58, 59], we assumed that higher frictional forces—indicative of greater integrin engagement and traction—lead to increased PI3K activation rates.

This signaling pathway influences the actomyosin network in two ways. First, the force generated by myosin motors, *ξ*, is made linearly dependent on the local concentration of active RhoA, such that regions with high RhoA exhibit stronger contractile forces, while areas with low RhoA show weaker myosin activity [12, 13, 14]. Second, the tension-free actin polymerization velocity at the leading edges of the cell, *v*^*p*^, is assumed to vary linearly with Rac1 density, meaning that actin polymerization is enhanced in regions where Rac1 accumulates [15, 16, 17, 18].

We then repeated the simulations described previously, now incorporating these additional biochemical interactions. The model reveals dynamic changes in Rac1 and RhoA levels, with both polarizing toward the friction gradient (see Fig. 1h-i). As a consequence, the retrograde flow becomes further polarized (Fig. 1j). We have assumed a sigmoidal-like response of PI3K activation due to friction, which saturates at high friction values. Hence, once the cell reaches region of high friction, the activation of PI3K by integrins becomes uniform from front to back, and therefore the Rac1 and RhoA (Fig. 1h-i). Altogether, the enhanced model predicts a higher migration speed compared to a cell influenced by a friction gradient alone (Fig. 1f).

Next, we extend this successful durotactic model to examine the interplay between electrical and mechanical cues and how they compete to direct cell migration (Fig. 2a). Specifically, our goal is to determine which cue—mechanical or electrical—is more effective in guiding cell migration. During electrotaxis, the electromigration of CMPs, from both electrophoretic and electroosmotic effects, leads to a polarized and enhanced activation of GTPases [70, 71, 34, 72, 73, 74, 37]. Therefore, both durotaxis and electrotaxis rely on the polarization of intracellular signaling pathways that regulate the cell’s motility machinery (Fig. 2a-b). To address this competitive behavior, we integrated the GTPase-based durotactic model described above, which is driven by mechanical stimulation, with our previously developed electrotaxis model (see SI and [69] for details), also employing the same GTPase signaling framework. Among the various possible downstream activation by CMPs, we first assume activation of PI3K by the electrically polarized CMPs [73, 72].

**Figure 2.**
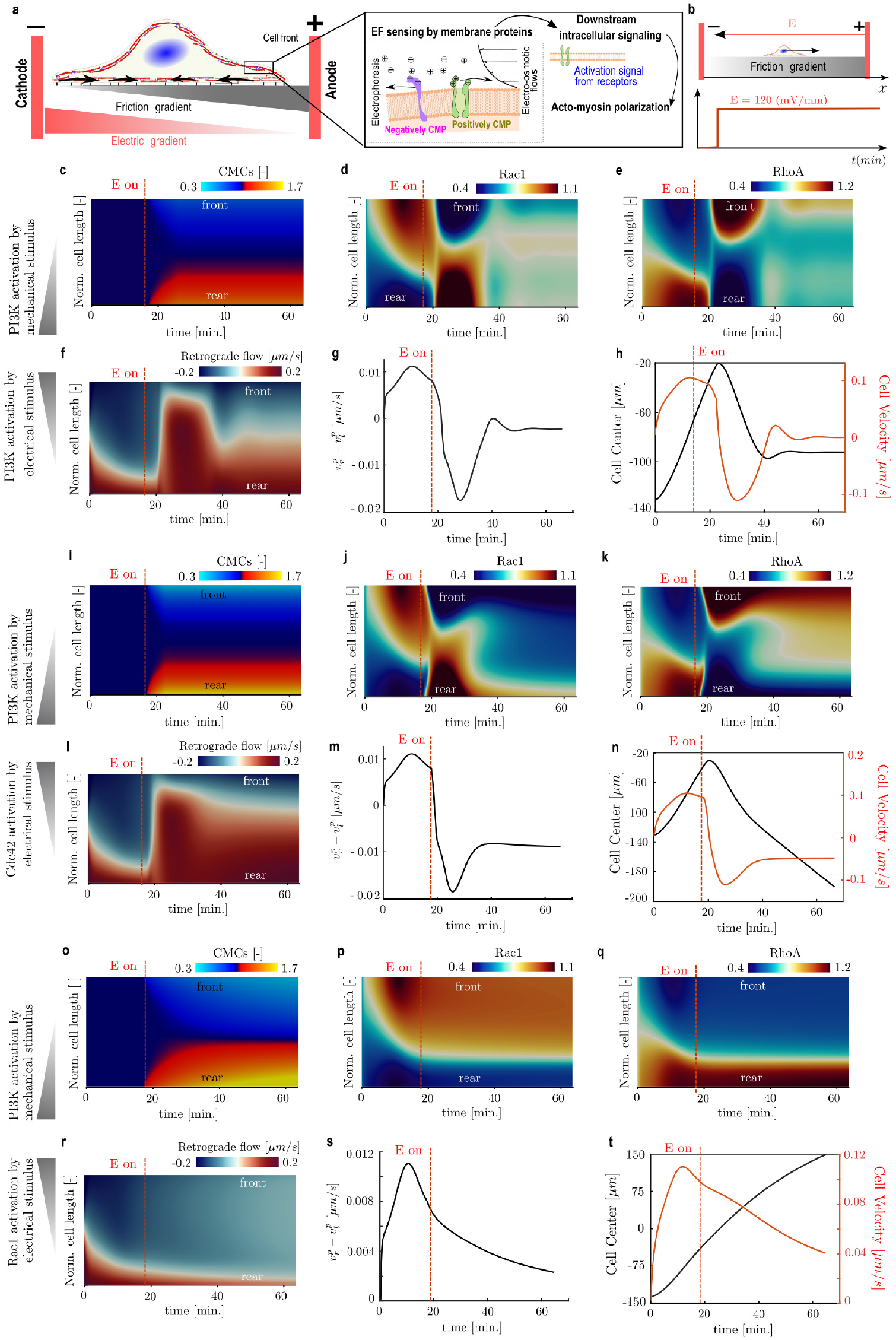
Competition of durotaxis and electrotaxis in single cell migration. (a) Opposing cues and sketch of the main mechanisms involved in electrotaxis: activation of CMPs, downstream signaling, and actomyosin polarization. (b) Sketch of the opposing activation of the electric field as a cell migrates along a friction gradient. (c-h) Polarization of CMPs induces a downstream activation of PI3K and, consequently, of the rho GTPases. (c-f) Kymograph of model variables along the normalized cell length (Y axis) over time (X axis). The kymographs show CMPs (c), Rac1 (d), RhoA (e), and the retrograde flow (f). (g) Difference in the polymerization velocity of actin against the cell membrane. (h) The position of the cell center (black lines) and migration velocity of the cell (red) is shown. (i-n) Polarization of CMPs induces a downstream activation of Rac1 and, consequently, of the rho GTPases. (i-l) Kymograph of model variables along the normalized cell length (Y axis) over time (X axis). The kymographs show CMPs (i), Rac1 (j), RhoA (k), and the retrograde flow (l). (m) Difference in the polymerization velocity of actin against the cell membrane. (n) The position of the cell center (black lines) and migration velocity of the cell (red) is shown. In all kymographs, we indicated the position of the front and rear of the cell. (o-t) Polarization of CMPs induces a downstream activation of Rac1 and, consequently, of the rho GTPases. (o-r) Kymograph of model variables along the normalized cell length (Y axis) over time (X axis). The kymographs show CMPs (o), Rac1 (p), RhoA (q), and the retrograde flow (r). (s) Difference in the polymerization velocity of actin against the cell membrane. (t) The position of the cell center (black lines) and migration velocity of the cell (red) is shown.

We apply the same stiffness gradient, position the cell center at −125 *μ*m, and allow the cell to migrate, as before. Thereafter, when the leading edge of the cell reaches *x* = −20 *μ*m, we activate an EF of 120 mV/*μ*m, with the cathode positioned at the rear of the migrating cell (Fig. 2b). Therefore, a stiffness gradient and an EF in the opposite direction are being applied. During this period, signals from the frictional gradient and CMP polarization toward the anode compete, leading to opposing stimuli for the activation of the signaling layer (see Fig. 1g) and, eventually, for the polarization of the cell.

Once the EF is applied, CMPs redistribute rapidly, on the order of seconds (Fig. 2c). There is an additional delay between EF activation and the subsequent change in migration direction, due to the turnover dynamics of PIPs and Rho GTPases (Fig. 2d–e). Specifically, RhoA-driven actomyosin flow (Fig. 2f) and Rac1-driven protrusive polymerization are modulated (Fig. 2g). As a result, we observe a reversal in migration velocity due to the change in migratory direction (Fig. 2h), indicating that electrotaxis can revert the migration along the stiffness gradient. For this specific set of model parameters, or cell-type, the migration velocity cancels after 30 min, when it reaches the steady state of stimulation, indicating that the effect of the opposed activation of PI3K compensates to each other (Fig. 2h).

There are also data indicating that activation by CMPs may directly stimulate the activation of Rac1 and Cdc42 [75, 76, 77, 78]. To analyze these possibilities, we first recompute the previous simulation with the direct activation of Cdc42 by CMPs instead of PI3K (Fig. 2i-n). Results are similar to the PI3K-based activation. However, we observe a re-polarization of Rac1 and RhoA (Fig. 2j-k), instead of the uniform distribution obtained for the PI3K-based activation, because the effect of the electric stimuli is larger than the PI3K-based activation. This polarization also creates a re-polarization of the retrograde flow (Fig. 2l), which is, in part, responsible for the reversal in migration direction (Fig. 2m). The cell reaches a steady velocity migration toward the cathode (Fig. 2n).

We also recompute the simulation with the activation of Rac1 by CMPs (Fig. 2o-t). We do not observe migration reversal but a reduction of migration speed (Fig. 2t) during the same period of time as in the previous simulation. After the initial polarization of Rac1 and RhoA (Fig. 2p-q), and consequently of the retrograde flow (Fig. 2r), we show a steady state. These results indicate that activation of Rac1 by the EF, opposed to the PI3K activated by a mechanical stimulus, is not enough to change the migration direction, and it just acts to reduce the migration capacity. Therefore, Rac1 seems to be a weaker effector for directed cell migration.

## Conclusions

Overall, our results show how the competition between electrical and mechanical stimuli in guiding cell migration arises from the activation of GTPase signaling through integrins or membrane proteins, and that the relative strength of these activation pathways ultimately determines which cue prevails. For the specific model parameters used in this work, our results suggest that PI3K is the most determining regulator in orchestrating the signaling state of GTPases in the cell and, therefore, symmetry breaking, polarity, and directional migration of the cell.

At this point, it became evident that determining whether mechanical or electrical stimulation dominates over each other is not a universal question but one that likely depends on the specific cell type. Although our model integrates the main mechanisms underlying durotaxis and electrotaxis, it involves a large number of parameters that strongly determine the effect of each Rho GTPase over the others in a cell-specific manner. And, therefore, it cannot fully resolve this question without complementary experimental validation. Thus, a key outcome of our modeling effort is the identification, formulation, and validation of specific experimental strategies needed to address this issue in a cell-type-dependent manner. As we discussed, central to both types of stimuli is the GTPase signaling network, which serves as a convergence point for upstream cues and downstream effects [10, 11]. In this work, we have adopted a well-established signaling framework that captures the interactions between GTPases and their regulators [66, 67, 68]. However, variations in feedback strengths, activation-deactivation kinetics, and receptor coupling are expected across different cell types. Fortunately, these aspects are readily available through mathematical modeling, and our model can provide accurate predictions to test at minimal cost.

The first step in dissecting durotactic and electrotactic behavior should thus focus on characterizing the signaling profile of the cell under study, specifically on integrin forms and all membrane components responsible for the GTPases signaling [79, 10, 80]. Indeed, different integrin subunits have been demonstrated significant regulatory roles in basal motility of cells, durotaxis and haptotaxis, and electrotaxis, albeit separately [81, 82, 83]. Selective expression of integrin subunits and membrane components could determine the migration direction of cells in an electric field, mediated by PI3 kinase signaling [83]. Thus, to test directly the role of integrin subunits and membrane proteins that regulate PI3K and GTPases to mechanical and electrical cues, one can manipulate the expression of these key membrane proteins with tunable CRISPR-interference (CRISPRi) or photoactivatable cas-13 [84]. This would allow for a precise spatio-temporal perturbation while assessing semi-quantitatively the contribution of each particular integrin subunit and membrane protein. These datasets would help in the experimental design of targeted perturbations and identify the upstream components most relevant for a given context. In the context of electrotaxis, preliminary screening of CMPs [85, 69] and integrins [83] has been attempted, but must be more closely investigated.

One global approach to understand CMPs would be to the tracking of membrane microdomains/rafts, which are composed of specific lipid environments and transmembrane proteins such as receptors and ion channels. Even if we do not know what proteins comprise these rafts at first, the indication that the lipid rafts have a compound nominal charge, because they have many identical proteins together, means that the raft can easily be “more positive” or “negative” and thus be responsive to the electric field [86]. Moreover, rafts have specific lipids like cholesterol that are uncharged, which can make the membrane more fluidlike because there’s less electrostatic interaction with lipid tails and thus less compaction. Markers for lipid rafts like FilipinIII and CTxB can be used to understand whether polarization of CMPs is dependent on protein-lipid complexes or whether individual components are sensing the EF and responding through electrophoresis and electro-osmotic flows. Other candidate molecules, such as EGFR or VEGFR, could be labeled for single-molecule microscopy, to be tracked during EF activation and determine what CMPs respond and repolarize to an EF. Another approach would be to knock down electrosensors such as VSP1, Galvanin (TMEM154), and PTEN and study the effect on electrotaxis direction [36, 87, 88, 89]. To further test our models regarding PI3K, an optogenetic controlled PI3K can be used to control its activity and that way, create an opposing signal to either durotaxis or electrotaxis [90]. Another way to test how determinant PI3k is for polarity and onset of migration is to use cell lines depleted for PI3K or use inhibitors, and check whether directionality is maintained with the biasing cues.

One outstanding question is whether PIPs are passively displaced by membrane flows upon EF activation. Therefore, tracking PIP2 or PIP3 fluorescent markers during EF activation could reveal whether PIPs are physically drifted or synthesized de novo at emerging polarity sites, and potentially whether the EF directly mobilizes the enzymatic machinery responsible for PIP phosphorylation and dephosphorylation. Since GTPases have a complex affinity to bind PIP kinases, PIP2, or even require PIP2 to activate their effectors, GTPases and PIP dynamics should be tracked in parallel to understand these relationships at the onset of cell repolarization. Many of these experiments or experimental strategies have already been applied in chemotaxis, hence a great inspiration and set of toolboxes can be easily adapted from there [91]. Besides these ideas, if one truly aims to understand how multimodal cues work in additive, synergistic, or confronting manners, a key direction is to characterize the environment in which cells migrate. This will deeply benefit theory and experimental design as it will report on the native scenarios in which cells move. For example, one can design experiments in which orthogonal, parallel, or antiparallel mechanical and electrical gradients, similar to those presented above, are applied while monitoring the parameters described above.

In the case of mechanical inputs, traction force microscopy paired with controlled substrate stiffness or micropatterned substrates [82] can be used to quantify how specific integrins modulate GTPase activation dynamics. Furthermore, to determine what integrins and CMPs, and at what strength and timing, activate which layers of the GTPase cascade, time-resolved imaging of integrin and CMPs clustering and associated GEF/GAP activity would be essential. This can be approached by live-cell FRET biosensing to monitor activities of Rac1, Cdc42 and Rho in real time during durotaxis/haptotaxis and upon the onset of an EF [92, 93], which can be combined with genetic perturbation (e.g., CRISPR-Cas9-mediated knockout or overexpression of specific integrins/CMPs). The onset of an EF can be controlled in direction and magnitude of the applied EF in an electrotaxis chamber, which allows direct test and validation of the modelling results as shown in both Figs 1 and 2. These experiments will directly compare the model dynamics of GTPase dynamics and the changes upon an EF being switched on.

Finally, how each Rho GTPase quantitatively influences actomyosin contractility has been much further investigated. For example, a common implementation consists of the use of myosin II fluorescent reporters (e.g., MRLC-GFP), in conjunction with pharmacological modulation of individual GTPases (using inhibitors or constitutively active mutants) and time-lapse confocal microscopy [94, 95]. Quantitative image analysis would then reveal how each GTPase contributes to cytoskeletal contractility in a stimulus-dependent manner. More importantly, and going back to the idea of synergy, addition or competition, the contribution of piezo electric membrane channels, such as TRPs or Piezos, which can rapidly deliver calcium influxes due to changes in membrane voltage and stretch, will also contribute to dissecting the molecular mechanism by which cells sense and translate electromechanical stimuli into a cellular pattern of actomyosin contractility.

Together, these experimental strategies would provide the necessary resolution to determine, for any given cell, which stimulus exerts dominant control over directional migration. Integrating these cues could lead to the development of efficient and controllable strategies for guiding cell migration, with significant implications for limiting cancer invasion, promoting wound healing, and engineering structured tissues on demand.

## Acknowledgments

PS acknowledges the support of the Spanish Ministry of Science and Innovation (Grant PID2022-142178NB-I00 funded by MCIN/ AEI/10.13039/501100011033/ FEDER, UE). S.K. was supported by the Spanish Ministry of Economy and Competitiveness (Grant number: PRE2020-095851). Work at the Barriga lab is supported by the European Research Council Starting Grant (ERC-StG) under the European Union’s Horizon 2020 research and innovation programme, Grant agreement No. 950254; the European Molecular Biology Organization (EMBO) Young Investigator Programme, Project No. 5248; and by the Deutsche Forschungsgemeinschaft (DFG, German Research Foundation) under Germany’s Excellence Strategy (EXC 2068, 390729961, Cluster of Excellence Physics of Life of TU Dresden).

