## Supplementary material for "Computational insights on the competition between electrotaxis and durotaxis": SI

January 23, 2026

### 1 Model description

We briefly outline the construction of the model, which largely follows the approach presented in our previous publications [1, 2, 3]. In essence, we consider a one-dimensional computational model. The cellular domain, $\Omega$, is represented as a continuous segment with moving boundaries $x(t) \in [l_l(t), l_r(t)]$, where $l_l(t)$ and $l_r(t)$ denote the positions of the left and right edges of the cell, respectively. Accordingly, the cell length is defined as $L(t) = |l_l(t) - l_r(t)|$. The velocities of the cell boundaries are given by $\dot{l}_l(t)$ and $\dot{l}_r(t)$, as described below. The displacement of the cell ends is governed by the velocity of the contractile actomyosin network, or retrograde flow, as well as by the actin polymerization velocity at the cathodal and anodal fronts, which ultimately determine the direction and speed of cell migration.

To model the cascade of events leading to mechanotaxis and electrotaxis, we consider the primary cellular sensors involved, integrins and Charged Membrane Proteins (CMPs), respectively, as described below.

#### 1.1 Cell adhesion based on the clutch model

We adopt clutch models to describe how cell adhesion responds to the mechanical properties of the extracellular matrix (ECM). The mathematical formulation, summarized below, follows previous work [4, 5]. In this framework, each molecular clutch connects the ECM on one side to the adaptor protein talin on the other. Talin, in turn, links to actin filaments, which are pulled by myosin motors. The actomyosin network generates contractile forces that act through these clutches at adhesion sites.

The velocity of the actin filaments, denoted by $\hat{v}$, follows a force–velocity relationship,

$$\hat{v} = v_u \left(1 - \frac{F}{F_{stall}}\right),$$

where $v_u$ is the unloaded actin flow velocity. The stall force of the myosin motors, $F_{stall}$, is defined as $F_{stall} = n_m F_m$, with $F_m$ representing the force required to stall a single myosin motor and $n_m$ the number of motors present. The reaction force exerted by the substrate is given by

$$F = \sum_{i=1}^{n_{eng}} F_{c,i},$$

where $n_{eng}$ denotes the number of engaged clutches and $F_{c,i}$ is the force transmitted through the $i$-th clutch.

The force transmitted by each clutch is calculated as

$$F_{c,i} = \kappa_c (z_{c,i} - z_{sub}),$$

where $\kappa_c$ is the stiffness of an individual molecular clutch [5], $z_{c,i}$ is the displacement of the $i$-th clutch, and $z_{sub}$ is the substrate displacement. Once the force on each clutch, $F_{c,i}$, is determined, we compute the corresponding binding and unbinding dynamics. The attachment of the $n_c$ clutches to the ECM, representing integrin–ECM interactions, is governed by catch-bond kinetics [8, 7]. Clutches bind at a constant rate $k_{on}$ and unbind at a force-dependent dissociation rate $k_{off}$. Mechanosensitivity arises through talin unfolding, which promotes vinculin recruitment and integrin clustering [7]—a process known as adhesion reinforcement. For further details on catch-bond behavior, mechanosensitivity modeling, and parameter values, see [5]. Once the total force $F$ is computed, the cell traction is obtained as

$$P = \frac{F}{\pi a^2},$$

where $\pi a^2$ is the area of a circular adhesion complex.

To compute the dynamics of the clutch model, we perform Monte Carlo (MC) simulations involving repeated stochastic sampling of clutch engagement and disengagement events. These cycles of adhesion formation and rupture are simulated using a constant time step $\Delta t = 0.005$ s over a total duration of $t_f = 100$ s, as validated in [5]. The displacement of engaged clutches during each time step is given by $\Delta z = \hat{v}\Delta t$. The substrate displacement at the next time step, $z^{n+1}$, is obtained by solving the force balance between the $n_{eng}$ engaged clutches and the substrate, which then allows for updating the total force $F$. Simulation outputs are averaged over time to capture the behavior under specific ECM mechanical properties.

Finally, we assume that the effective friction between the cell and the ECM is given by [6, 2]

$$\eta = \frac{P}{\hat{v}},$$

This relationship implies that friction varies with ECM stiffness in a manner similar to the changes in actin velocity and traction force. Under this assumption, strong and stable adhesions—reflected by higher traction forces—correspond to increased friction, whereas weak adhesions produce lower friction.

#### 1.2 Electromotility of charged membrane components

The electro-motility of CMPs arises from the interplay between two opposing forces. When a cell is immersed in an electrolyte solution, its negatively charged membrane attracts nearby positive ions, forming a structured region known as the electrical double layer (EDL). Upon the application of an electric field (EF), this EDL is also subjected to the field, generating a fluid movement known as electro-osmotic flow (EOF). CMPs embedded in the cell membrane are dragged by this EOF, enabling their migration along the membrane surface. In addition to being influenced by the global EOF, the charged nature of CMPs causes them to experience local EOF as well as electrostatic interactions. These combined effects constitute electrophoresis, which leads to an asymmetric redistribution of CMPs along the cell membrane [9, 10, 11].

To quantify the rate and magnitude of this redistribution—which ultimately drives the downstream polarization of intracellular signaling pathways—we adopted previously established mathematical models [9, 10, 11, 3]. In summary, the total electro-motility velocity can be expressed as

$$v_e = \frac{\epsilon_r \epsilon_0 E(\zeta_1 - \zeta_2)}{\eta_m}, \tag{1}$$

where $\epsilon_r$ is the relative dielectric constant of the surrounding medium, $\epsilon_0$ is the vacuum permittivity, $E$ is the strength of the applied EF, and $\eta_m$ is the viscosity of the cell membrane. The terms $\zeta_1$ and $\zeta_2$ represent the $\zeta$-potentials of the CMPs and the cell surface, respectively.

To model the spatiotemporal distribution of positively and negatively charged CMPs, denoted by $\rho_\pm$, we use a convection–diffusion equation:

$$\partial_t \rho_\pm + \partial_x \left( v_e \rho_\pm - D_M \partial_x \rho_\pm \right) = 0, \tag{2}$$

where $D_M$ denotes the diffusion coefficient of CMPs. We impose zero Neumann boundary conditions at both ends, assuming that CMPs neither enter nor leave the cell membrane. The initial condition is set as a normalized uniform distribution, $\rho_\pm(x,0) = 1$.

### 1.3 Polarization of signaling cues by a GTPases model

To incorporate the complex interactions between Rho GTPase, specifically active and inactive forms of Rac1, Cdc42, and RhoA, as well as key phosphoinositides such us PIP ($P_1$), PIP2 ($P_2$), and PIP3 ($P_3$), we take a model described in our previous publication on electrotaxis [3], which was based on several previously proposed models [12, 13, 14, 15, 16, 17].

In summary, the active forms of Rac, Cdc42, and Rho are, respectively:

$$\frac{\partial R}{\partial t} = (I_R + \alpha C)\left((1-f_2) + f_2\frac{P_3}{P_{3b}}\right)\frac{R_i}{R_{tot}} - d_R R + D_m \partial_x^2 R, \tag{3}$$

$$\frac{\partial C}{\partial t} = \left(\frac{I_C}{(1+(\frac{\rho}{a_1})^n}\right)\left((1-f_1) + f_1\frac{P_3}{P_{3b}}\right)\frac{C_i}{C_{tot}} - d_C C + D_m \partial_x^2 C \text{ and} \tag{4}$$

$$\frac{\partial \rho}{\partial t} = (\frac{(I_\rho + \beta R)}{(1+(\frac{C}{a_2})^n})\frac{\rho_i}{\rho_{tot}} - d_\rho \rho + D_m \partial_x^2 \rho, \tag{5}$$

respectively. The C$_{tot}$, R$_{tot}$, $\rho_{tot}$ are the total concentrations of the Cdc42, Rac and Rho. $I_c, I_R$ and $I_\rho$ are activation rates, which we describe below. The values $a_1$ and $a_2$ are the Rho and Cdc42 concentrations that elicit a half-maximal drop of Cdc42 and Rho activation, respectively. The value $\alpha$ determines the rate of Cdc42-enhanced activation of Rac, and $\beta$ determines the rate of Rac-enhanced Rho activation. $P_{3b}$ is the baseline concentration of PIP3 found in a resting cell. $d_R$, $d_C$, and $d_\rho$ are the decay rates of activated Rac1, Cdc42 and Rho, respectively. $D_m$ is the diffusion constant of the active form. $f_1$ and $f_2$ are feedback from PIP$_3$ to Cdc42 and Rac, respectively, which we assume differs from the original model [13], and they are bounded such that $0 \leq f_1, f_2, \leq 1$. This allows us to analyze different feedback strengths from PIP3 to Rac1 and Cdc42.

$R_i, C_i, \rho_i$ are the concentrations of the respective inactive forms, which satisfy the equations:

$$\frac{\partial R_i}{\partial t} = -(I_R + \alpha C)\left((1-f_2) + f_2\frac{P_3}{P_{3b}}\right)\frac{R_i}{R_{tot}} + d_R R + D_{mc} \partial_x^2 R, \tag{6}$$

$$\frac{\partial C_i}{\partial t} = -\left(\frac{I_C}{(1+(\frac{\rho}{a_1})^n}\right)\left((1-f_1) + f_1\frac{P_3}{P_{3b}}\right)\frac{C_i}{C_{tot}} + d_C C + D_{mc} \partial_x^2 C \text{ and} \tag{7}$$

$$\frac{\partial \rho_i}{\partial t} = -\frac{(I_\rho + \beta R)}{(1+(\frac{C}{a_2})^n}\frac{\rho_i}{\rho_{tot}} + d_\rho \rho + D_{mc} \partial_x^2 \rho. \tag{8}$$

D$_{mc}$ is the diffusion coefficient of inactive Rho-proteins, which averages the respective diffusion rates of inactive GTPase forms by the time spent on the membrane versus the cytosol and diffuses much faster than the active forms (D$_m \ll$ D$_{mc}$).

Interactions between GTPases and PIPs are included in coupling terms between these sets of PDEs. The interaction between phosphoinositides, PIP ($P_1$), PIP2 ($P_2$), and PIP3 ($P_3$), and between phosphoinositide kinases, are also modeled by a set of similar PDEs with reaction kinetics as

$$\partial_t P_1 - D_p \partial_x^2 P_1 = I_{P1} - \delta_{P1} P_1 + k_{21} P_2 - f_{PI5K} P_1 \tag{9}$$

$$\partial_t P_2 - D_p \partial_x^2 P_2 = -k_{21} P_2 + f_{PI5K} P_1 - f_{PI3K} P_2 + f_{PTEN} P_3 \tag{10}$$

$$\partial_t P_3 - D_p \partial_x^2 P_3 = f_{PI3K} P_2 - f_{PTEN} P_3 \tag{11}$$

where it is assumed that the conversion to PIP occurs at a constant rate, $I_{P1}$, and that all PIs diffuse in the membrane at a uniform rate $D_p$. $\delta_{P1}$ is the PIP$_1$ decay rate and $k_{21}$ is the PIP$_2$ to PIP$_1$ conversion rate. The model also describes the fact that Rac enhances the conversion of PIP to PIP2 (via PI5K), of

PIP2 to PIP3 (via PI3K), and that Rho enhances the conversion of PIP3 to PIP2 via PTEN. The feedbacks are incorporated through the functions

$$f_{PI5K} = \frac{k_{pi5k}}{2}(1 + \frac{R}{R_b}), \tag{12}$$

$$f_{PI3K} = \frac{k_{pi3k}}{2}(1 + \frac{R}{R_b}), \text{ and} \tag{13}$$

$$f_{PTEN} = \frac{k_{pten}}{2}(1 + \frac{\rho}{\rho_b}), \tag{14}$$

where $k_{PI3K}$ is the $PIP_2$ to $PIP_3$ baseline conversion rate, $k_{PI5K}$ is the $PIP_1$ to $PIP_2$ baseline conversion rate and $k_{PTEN}$ is the $PIP_3$ to $PIP_2$ baseline conversion rate. Whenever we modify $k_{pi3k}$, we also simultaneously adjust $k_{pten}$, as they are the forward and backward rates of the same reaction. This is done with $\rho_-$ and the same strength factor $S_P$. $R_b$, $C_b$ and $\rho_b$ are typical levels of active Rac, Cdc42 and Rho respectively.

We modified this model to incorporate stimuli arising from integrins and CMPs. To do so, we assume that the activation rates of the relevant signaling components depend on the magnitude of CMP accumulation or integrin activation. First, we assume that integrin engagement activates PI3K according to

$$\dot{k}_{pi3k} = \lambda(S(\eta) - k_{pi3k}) \text{ with } S(\eta) = \frac{1}{1 + e^{-10(\eta - 0.6)}} \tag{15}$$

an activation function that depends on the friction or, in other words, on the activity of cell adhesion following a sigmoidal function, where $\lambda$ is a given relaxation rate that we fix to $\lambda = 0.5$ to describe a slow biochemical adaptation. The slow adaptation accounts for chemical activation cascades, PI3K recruitment time to the membrane, and possible buffering events.

Regarding activation of Rho GTPases by polarized CMPs, we assume a linear relationship of the form

$$I_C = \rho_\pm I_C^*, \qquad I_R = \rho_\pm I_R^*, \qquad k_{pi3k} = \rho_\pm k_{pi3k}^*, \tag{16}$$

for Cdc42, Rac1, and PI3K activation, respectively. Here, the activation scales with the baseline conversion rates $I_C^*$, $I_R^*$, and $k_{pi3k}^*$ defined in the base model [13].

These modifications represent modeling assumptions that may require validation or refinement for specific cell types.

### 1.4 Mechanical model of the actomyosin network

To describe the mechanics of the actomyosin network, we adopt an active gel model [18, 1], formulated as

$$\partial_x \sigma = \eta v^F \quad \text{in } \Omega. \tag{17}$$

Here, inertial terms are neglected, and the internal stress $\sigma$ is defined through a constitutive relation as

$$\sigma = -2\mu\, \partial_x v^F + \zeta\, \rho\, \rho^F \rho^M,$$

which accounts for both the viscous resistance of the actin cortex and the active contractility generated by the actomyosin system. In this expression, $\rho^F$ and $\rho^M$ represent the concentrations of F-actin and bound myosin motors, respectively, as detailed in the next section. The velocity $v^F$ corresponds to the actin network (or retrograde flow) measured in the laboratory frame. The parameter $\mu$ denotes the shear viscosity of the network, while $\zeta$ characterizes the active contractile stress generated by myosin motors. This contractility is further modulated by the local concentration of RhoA, $R$, thereby enhancing contractility in regions with elevated RhoA activity. Zero Neumann boundary conditions are imposed at both ends of the domain. The right-hand-side term in Eq. (17) represents the frictional interaction between the moving cortex and its surrounding environment. We assume this friction to be linearly proportional to the cortex velocity, with coefficient $\eta$ [1, 2], as defined above.

### 1.5 Kinetics of the cell fronts

The polymerization velocity of actin filaments at the cell front is biochemically regulated by the activity of Rac1 and Cdc42, which modulate Arp2/3 complex activation and filament nucleation [19]. In the absence of opposing forces, the polymerization velocity is defined as

$$v_0^p = k_{on}\, \delta\, R,$$

where $k_{on}$ represents the actin assembly rate at the leading edge, and $\delta$ denotes the size of a single actin monomer. Because $v_0^p$ depends on the concentration of active Rac1, $R$, actin protrusion is enhanced at the cell front, where Rac1 is abundant, and inhibited at the rear.

Polymerization is physically opposed by the membrane tension, $\tau(L(t))$. We assume that membrane tension follows a linear relationship given by

$$\tau = -k\left(L(t) - L_0\right),$$

where $k$ denotes the membrane stiffness and $L_0$ is the initial cell length.

To describe the dependence of polymerization on membrane tension, we adopt a well-established model introduced by [20], in which the polymerization velocity decreases with increasing tension according to

$$v^p = v_0^p \left[1 - \frac{\tau(L(t))}{\tau_{stall}}\right]^{\gamma}, \tag{18}$$

where $\tau_{stall}$ represents the membrane tension required to stall polymerization, and $\gamma$ determines how sharply the velocity decays with increasing tension.

Given both the retrograde actin flow velocity, $v$, and the actin polymerization velocity, $v^p$, the motion of the cell boundaries is obtained as a combination of inward flow and outward protrusion. Specifically, the velocities of the front and rear edges are defined as

$$\dot{l}_f(t) = v_f + v_f^p, \quad \text{and} \quad \dot{l}_b(t) = v_b + v_b^p,$$

respectively. The net migration velocity of the cell is then given by

$$v = \frac{\dot{l}_f(t) + \dot{l}_b(t)}{2}. \tag{19}$$

### 1.6 Transport of the actomyosin network

Finally, to model the density dynamics of the actomyosin network, we employ a coupled convection–diffusion framework to describe the spatiotemporal evolution of filamentous (F-actin), $\rho^F(x,t)$, and globular (G-actin), $\rho^G(x,t)$, actin species within the cell (see [1] and references therein):

$$\partial_t \rho^F + \partial_x\left(w\rho^F - D^F \partial_x \rho^F\right) = k_p \rho^G - k_d \rho^F, \tag{20}$$

$$\partial_t \rho^G - \partial_x\left(D^G \partial_x \rho^G\right) = k_d \rho^F - k_p \rho^G. \tag{21}$$

Here, $D^F$ and $D^G$ denote the diffusion coefficients of F-actin and G-actin, respectively, while $k_p$ and $k_d$ are the polymerization and depolymerization rates. Because G-actin diffuses much more rapidly than it is advected by flow, convection is neglected in its equation. The retrograde flow relative to the cell frame is defined as $w = v^F - v$. Zero-flux boundary conditions are imposed at both cell edges, assuming that neither form of actin crosses the membrane boundary.

In addition, we consider a two-species model for myosin motors: a bound form, $\rho^M$, attached to the F-actin network, and an unbound form, $\rho^m$. Their redistribution is similarly governed by convection–diffusion dynamics:

$$\partial_t \rho^M + \partial_x\left(w\rho^M - D^M \partial_x \rho^M\right) = k_b^M \rho^m - k_u^m \rho^M, \tag{22}$$

$$\partial_t \rho^m - D^m \partial_x^2 \rho^m = k_u^m \rho^M - k_b^M \rho^m. \tag{23}$$

Here, $D^M$ and $D^m$ denote the diffusion coefficients of bound and unbound myosin, respectively, while $k_b^M$ and $k_u^m$ represent the binding and unbinding rates of myosin to F-actin. As with G-actin, the diffusivity of unbound myosin is assumed to dominate, allowing convection to be neglected in Eq. (23). Zero-flux boundary conditions are again applied at both ends of the domain to ensure that neither form of myosin enters nor leaves the cell.

### 2 Numerical solution of the problem and model parameters

We numerically solve all coupled PDE systems using a staggered approach. Spatial discretization is performed using the Finite Element Method (FEM), while temporal discretization of the parabolic equations is carried out using the implicit, second-order Crank–Nicolson scheme [21]. Nonlinear equations are solved iteratively via the Newton–Raphson method. Unless otherwise specified in the main text, all simulation parameters are adopted from previously published studies [1, 2, 3].
